# A Multi-Niche Microvascularized Human Bone-Marrow-on-a-Chip

**DOI:** 10.1101/2019.12.15.876813

**Authors:** Michael R. Nelson, Delta Ghoshal, Joscelyn C. Mejías, David Frey Rubio, Emily Keith, Krishnendu Roy

## Abstract

The human bone marrow (hBM) is a complex organ critical for hematopoietic and immune homeostasis, and where many cancers metastasize. Yet, understanding the fundamental biology of the hBM in health and diseases remain difficult due to complexity of studying or manipulating the BM in humans. Accurate *in vitro* models of the hBM microenvironment are critical to further our understanding of the BM niche and advancing new clinical interventions. Although, *in vitro* culture models that recapitulate some key components of the BM niche have been reported, there are no examples of a fully human, *in vitro*, organoid platform that incorporates the various niches of the hBM - specifically the endosteal, central marrow, and perivascular niches – thus limiting their physiological relevance. Here we report an hBM-on-a-chip that incorporates these three niches in a single micro-physiological device. Osteogenic differentiation of hMSCs produced robust mineralization on the PDMS surface (“bone layer”) and subsequent seeding of endothelial cells and hMSCs in a hydrogel network (“central marrow”) created an interconnected vascular network (“perivascular niche”) on top. We show that this multi-niche hBM accurately mimics the ECM composition, allows hematopoietic progenitor cell proliferation and migration, and is affected by radiation. A key finding is that the endosteal niche significantly contributes to hBM physiology. Taken together, this multi-niche micro-physiological system opens up new opportunities in hBM research and therapeutics development, and can be used to better understand hBM physiology, normal and impaired hematopoiesis, and hBM pathologies, including cancer metastasis, multiple myelomas, and BM failures.

## Introduction

Hematopoietic stem cells (HSCs) reside and self-renew in the bone marrow (BM) throughout adulthood, where multipotency is maintained and hematopoietic progenitor cells (HPCs) differentiate to maintain hematopoietic homeostasis (*1*). The microenvironment that maintains HSC potency and regulates differentiation of HPCs, i.e. the HSC niche, is characterized by BM stromal cells, extracellular matrix (ECM), and biochemical and physical signals (*2–4*). The HSC niche can be disrupted, naturally with age, with radiation or chemotherapies, or by primary and metastatic malignancies in the BM, and also through the mobilization of HSCs for apheresis. Novel, human, and potentially patient specific, models of this microenvironment are critical to advancing our understanding of the BM niche, develop new BM directed therapeutics, and evaluate the effects and predict the success (or failure) of clinical interventions (*5*).

Our understanding of the location and composition of the BM microenvironment has been changing over the last two decades. Early research had indicated that HSCs resided in a hypoxic, endosteal niche, where potency was maintained (*6, 7*). However, recent findings have shown that multi-potent HSCs are perivascular and exist in a niche maintained by endothelial cells and perivascular stromal cells (*8–10*). Across these distinct microenvironments, a number of stromal cells, including osteoblasts, MSCs, endothelial cells, CXCL12-abundant reticular (CAR) cells, adipocytes, macrophages, and osteocytes have all been implicated in regulating HSC fate (*11–13*). As HSCs differentiate, hematopoietic progenitor cells (HPCs) occupy distinct microenvironments within the BM, where they differentiate into lymphoid and myeloid lineages (*14*).

The ability to replicate the juxtaposition and interaction of the endosteal and perivascular niches is important to understand the complexity of the human BM niche (Fig 1A). The endosteal niche is primarily constituted by osteoblasts (OBs) and osteoclasts. OBs express extracellular matrix (ECM) in the endosteal microenvironment (e.g., collagen I (Col I), fibronectin (FN), and osteopontin (OPN)). OBs also provide soluble and surface bound signals, like jagged-1 (JAG1) and stem cell factor (SCF), to HSCs and HPCs that regulate differentiation and potency. The endosteal niche was believed to be relatively hypoxic and this was thought to promote HSC maintenance of potency; however, recent studies have not found the endosteal niche to be hypoxic (*15*). In part, this is due to the high vascularization throughout the bone compartment. BM sinusoids permeate the bone cavity and serve as the connection between BM and peripheral tissues, allowing for the egress of progenitors of the hematopoietic system. Endothelial cells (ECs) comprise the BM sinusoids and are accompanied by mesenchymal stromal cells (MSCs) or pericytes in establishing the perivascular niche. Abundant levels of stromal derived factor 1 (SDF1 or CXCL12) and SCF are expressed in the perivascular space to establish the HSC niche and recent *in vivo* studies have observed HSCs to be resident in the perivascular niche (*16–18*).

**Fig. 1.**
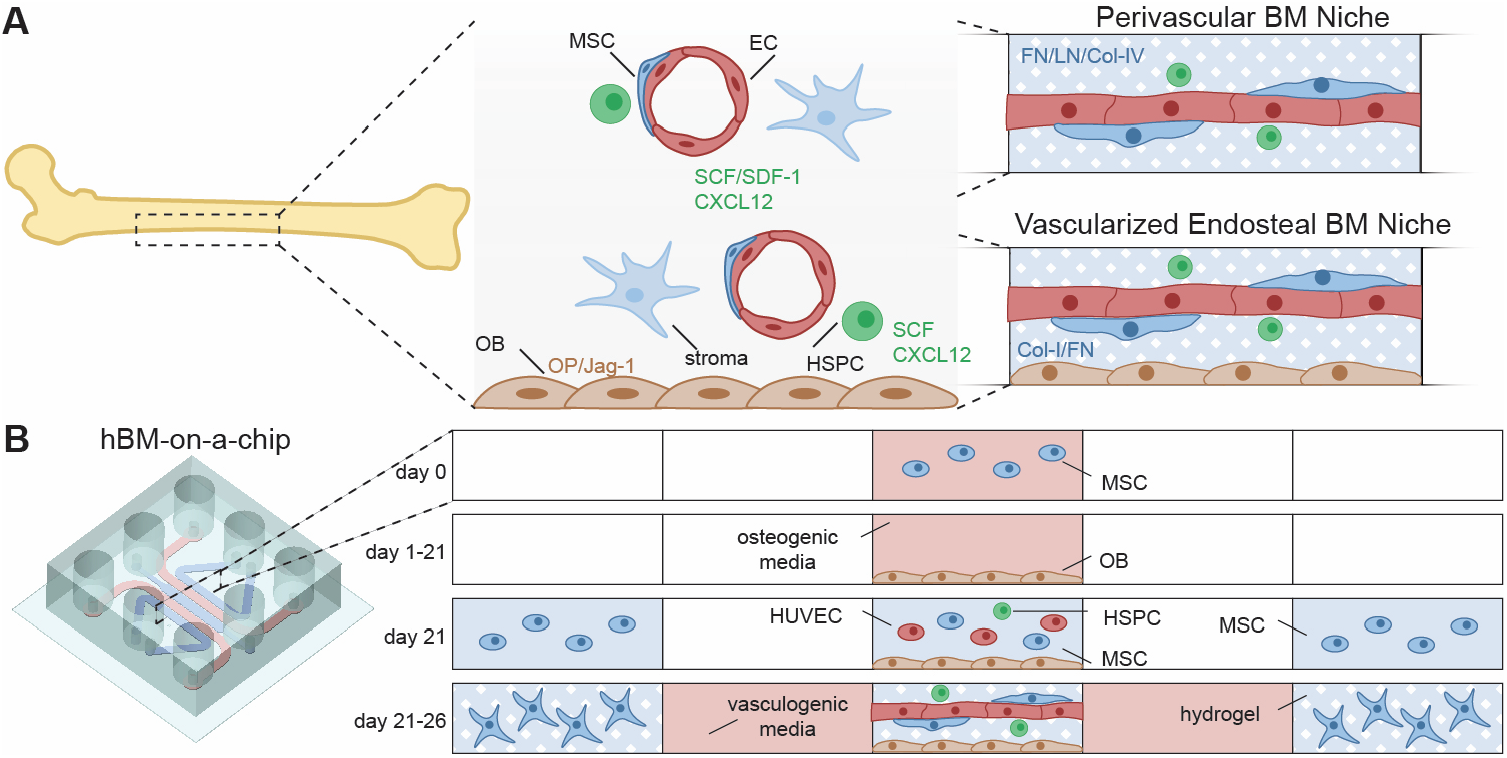
Schematic of human bone marrow-on-a-chip (hBM-on-a-chip). (A) The hBM-on-a-chip can recapitulate both the central perivascular BM niche (without OBs) and the vascularized endosteal BM niche (with OBs) that are found in the cavities of long bones. MSC = mesenchymal or marrow stromal cells, including pericytes; OB = osteoblasts and mineralized bone-like tissue layer; stromal cells = other cells of the BM stroma including CXCL12-abundant reticular cells (CAR), matured hematopoietic cells, and adipose cells. Note the area between vasculatures represent the central marrow region. FN = Fibronectin; LN = Laminin, Col-I and Col-IV = Collagen I and Collagen IV; OP = Osteopontin; Jag-1 = Jagged 1. (B) A 5-channel PDMS microfluidic device was fabricated using standard soft lithography techniques. MSCs are first differentiated for 21 days in the central channel of the device to form an endosteal layer, then HUVECs, MSCs, and HSPCs are loaded on top of the endosteal layer and vasculogenesis occurs over 5 days to form the hBM-on-a-chip.

The HSC niche has been a subject of study for decades and there have been many efforts to recapitulate aspects of the niche *in vitro*. Various material approaches and simple co-culture platforms in both 2D and 3D have been reported to increase HSC proliferation or maintain HSC potency during *in vitro* culture (*19–21*). Culture systems have been developed that mimic specific cytokine (*22*) or ECM (*23*) environments that HSCs experience in the BM niche *in vivo*. More recently, *ex vivo* and *in vitro* microfluidic or on-chip devices have been designed for recreating the bone or BM microenvironment (*24–26*). These studies recapitulate many characteristics of bone marrow; however, they either require lengthy ectopic implantation in animal models, or do not recreate the vasculature of BM, or do not incorporate the multiple juxtaposed niches of the BM microenvironment (especially the endosteal niche with the central marrow an dperivascular niche together), and ultimately are neither fully human nor complete models of the hBM.

Here, we present a simple microfluidic human BM-on-a-chip (hBM-on-a-chip) that can be used to create a human MSC-derived endosteal surface overlaid with a microvascular hydrogel network with hMSCs representing the central marrow and perivascular niches - that mimic the basic multi-niche structure of the human BM (Fig. 1B). We demonstrate the use of this device to study HSC response to a variety of stimuli, such as radiation and clinical HSC mobilization agents. This device can also be used to study human BM physiology and pathologies, including cancer metastasis, multiple myelomas, and bone marrow failures.

## Results

### Design and Fabrication of a Multi-niche hBM-on-a-Chip

A five-channel device was designed and fabricated with a height of 150 μm and consists of one central “gel channel”, two media channels, and two outer gel channels (Fig. S1A). The device was designed similarly to recently published methods (*27, 28*), but with modifications that (A) promoted maintenance of the air-liquid interface between the central channel and adjacent media channels, and (B) allowed for loading of cells and hydrogel precursors into a previously wetted channel. Essential to this approach is the ability to wet the central channel during the differentiation of hMSCs while keeping the adjacent media channels dry throughout the process so that a second set of cells can be loaded within the central channel. To achieve this, the communication pores between the gel channels were narrowed (compared to similar, previously published devices) to 50 μm in width and the number of communication pores were limited to decrease the occurrence of media leaking from the central channel during osteogenesis. The theoretical difference between advancing pressure and burst pressure (Supplementary Methods) was calculated to be 28.5 cm H_2_O which allows for simple and easy loading of fluid into the device. However, extended culture of the devices in a humid environment results in wetting of interior, dry surfaces of PDMS that leads to “failure” of the devices. This is mitigated by thorough cleaning of the devices with 70% EtOH and subsequent drying at 65 °C and can produce devices that “survive” the 21-day differentiation at a rate >90% (Fig. S2B).

This approach requires the central channel to be accessible by both a small port (1 mm) for initial and subsequent loading of cells, and a larger media reservoir (4 mm) for extended culture of the hMSCs during osteogenic differentiation. During our initial studies, we approached this challenge by fabrication of a 3-layer PDMS device (Fig. S1D, E) consisting of (a) a PDMS coated glass coverslip, (b) a middle-feature layer, and (c) a reservoir layer. This fabrication method was appropriate for initial studies because it allowed for fabrication of individual devices while failure rates were high, and its modular composition permitted parallel iteration of parts. However, as the design became finalized, this fabrication method proved to be inefficient and time intensive. To improve efficiency of fabrication, we developed a simplified method that combined the feature layer and the reservoir layer through a two-step PDMS casting protocol where a 3D printed mold was used to create media reservoirs (Fig. S1F, G). This fabrication protocol substantially decreased device fabrication time and increased material efficiency.

To further improve on the design and standardization of hBM-on-a-chip, a microfluidic platform was developed to integrate the platform into a standard, well-plate format. Using previously published methods (*29–31*), the PDMS “device” layer was bonded directly to a commercially available, bottomless 96-well plate (Fig. S1H, I). The resulting array of 8 devices uses the wells of the plate as media reservoirs and as a window for imaging the central channel of the device (Fig. S1J, K). This process creates an array of devices that are consistently oriented within known well locations, making the platform easily transferable to automated imaging, or potentially, media handling instrumentation. An unexpected benefit of moving to this fabrication method was the increased “survival” of devices during the 21-day differentiation of MSCs. This was most likely because the PDMS portion of the construct is fixed to a rigid polystyrene frame, so there were no longer small deformations due to handling of the devices during loading and culture, resulting in a much lower frequency, less than 5%, of device “failure” (Fig. S2C).

### On-Chip Osteogenic differentiation of hMSCs Create Mineralized Endosteal Niche

To promote cell adhesion to the bottom surface of PDMS within the central channel of the device, microfluidic devices were coated with polydopamine (PDA) and collagen I(*32*), which was found to improve attachment of hMSCs when compared to collagen I or fibronectin alone (Fig. S2A). hMSCs were then seeded at a high density within the central channel of the device and differentiated with osteogenic media over a period of 21 days. Mineralization of matrix was observed by Alizarin red (Fig. 2A, B) and von Kossa (Fig. 2C, D) staining. Mineralization increased over the 21-day differentiation. At 21 days, 71±18% of the area of the device stained positive for Alizarin red and the normalized mean intensity of the stained device was 0.64±0.07. Similarly, at 21 days 91±3% of the area of the device stained positive for von Kossa and the normalized mean intensity was 0.60±0.06. In addition to mineralized matrix, the expression of specific cytokines and ECM components is of interest when creating a surrogate endosteal niche. After 21 days differentiation, the differentiated hMSCs expressed cytokines (SCF, CXCL12, JAG1) and ECM (FN, OPN) characteristics of the endosteal niche (Fig. 2E-I).

**Fig. 2.**
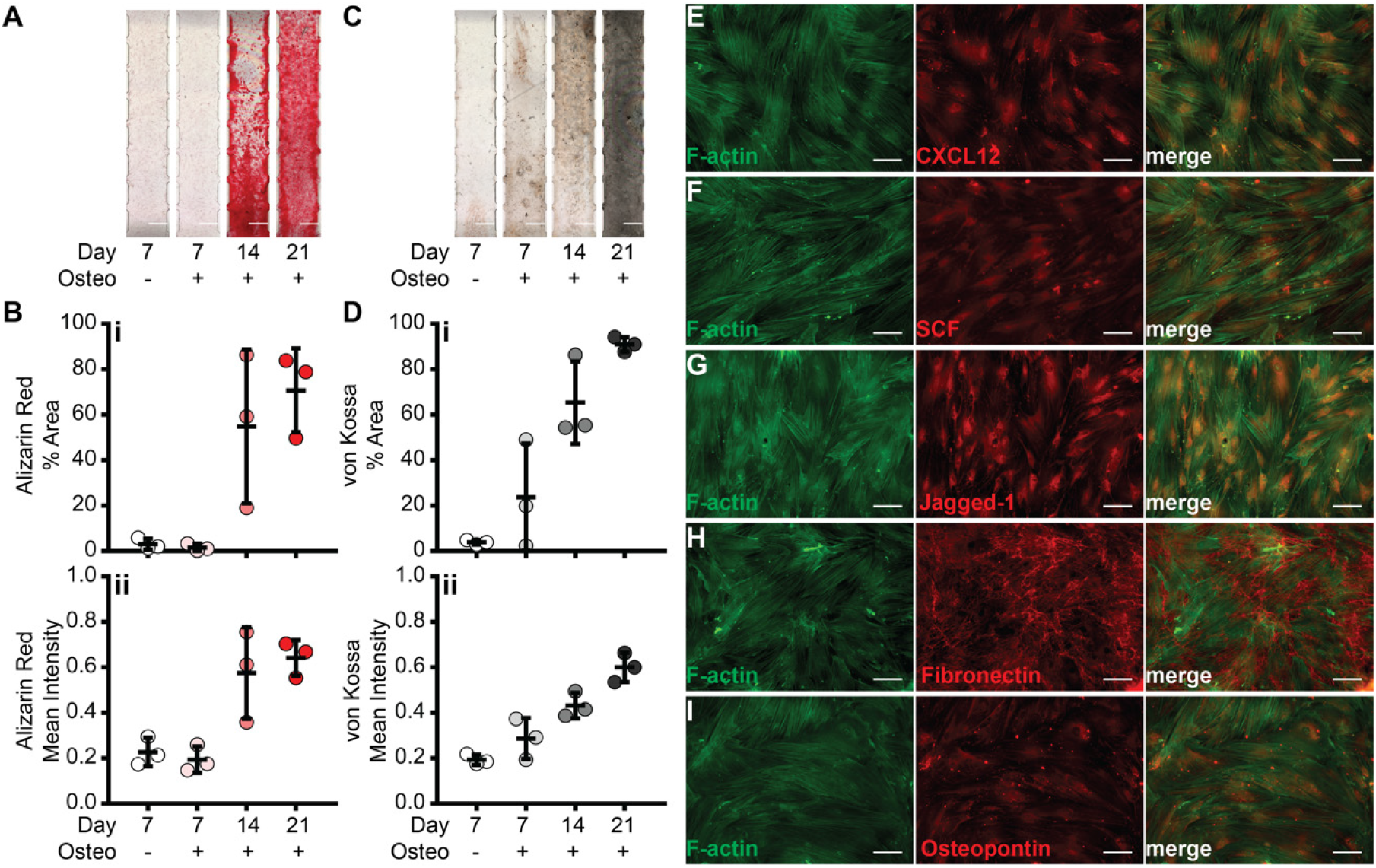
Formation of the endosteal niche. MSCs were differentiated for 21 days within the central channel of the hBM-on-a-chip after which mineralization was measured. Representative images of (A) alizarin red and (C) von Kossa staining. Scale bars: 500 μm. Quantification of (B) alizarin red and (D) von Kossa staining by (i) percent area and (ii) mean intensity. Data are plotted as mean ± SD (n = 3 devices). Immunofluorescence staining of endosteal cytokines (E) CXCL12, (F) SCF, and (G) jagged-1. Immunofluorescence staining of endosteal ECM (H) fibronectin, and (I) osteopontin, was observed using immunofluorescence staining. Scale bars: 100 μm.

### hMSCs co-cultured with Endothelial Cells in a Fibrin-Collagen Gel Allows Robust Generation of the Central Marrow and Perivascular Niche on the Endosteal Niche

In recent years, several groups have published methods for *in vitro* vasculogenesis in similar microfluidic platforms (*27, 28, 33, 34*). We found that perfusion of the vasculature was most consistent using a combination of hMSCs in laterally adjacent channels, supplementation with vascular endothelial growth factor (VEGF) and angiopoietin-1 (ANG-1), and encapsulation of endothelial cells in a fibrin/collagen co-gel (Fig. S3). Seeding HUVECs and hMSCs on top of the newly formed endosteal surface and culturing under these conditions for 5 days created vasculature in a reproducible manner.

### Effect of stromal cells on vasculogenesis

hBM-on-a-chip microfluidic devices were created with and without hMSCs and the differentiated endosteal layer (OBs) to measure the effect of stromal cells on vasculogenesis and cytokine secretion (Fig. 3A). We observed no significant difference in the vasculature area containing hMSCs (41.4±4.4%), OBs (44.1±2.9%), or both cell types (44.3±3.1%) when compared between groups or to devices containing ECs only (43.39±3.3%) (Fig. 3B). Similarly, no significant difference in the total length of the vasculature networks was observed between devices containing hMSCs (83.0±13.0 mm), OBs (92.9±12.2 mm), both hMSCs and OBs (92.1±10.8 mm), or neither (83.3±6.2 mm) (Fig. 3C). Although there was no significant difference, it is worth noting that the devices containing OBs exhibited vasculature that covered marginally less area and had a slightly increased total length of the network, which indicates a smaller average diameter of the newly formed vasculature compared to devices without OBs.

**Fig. 3.**
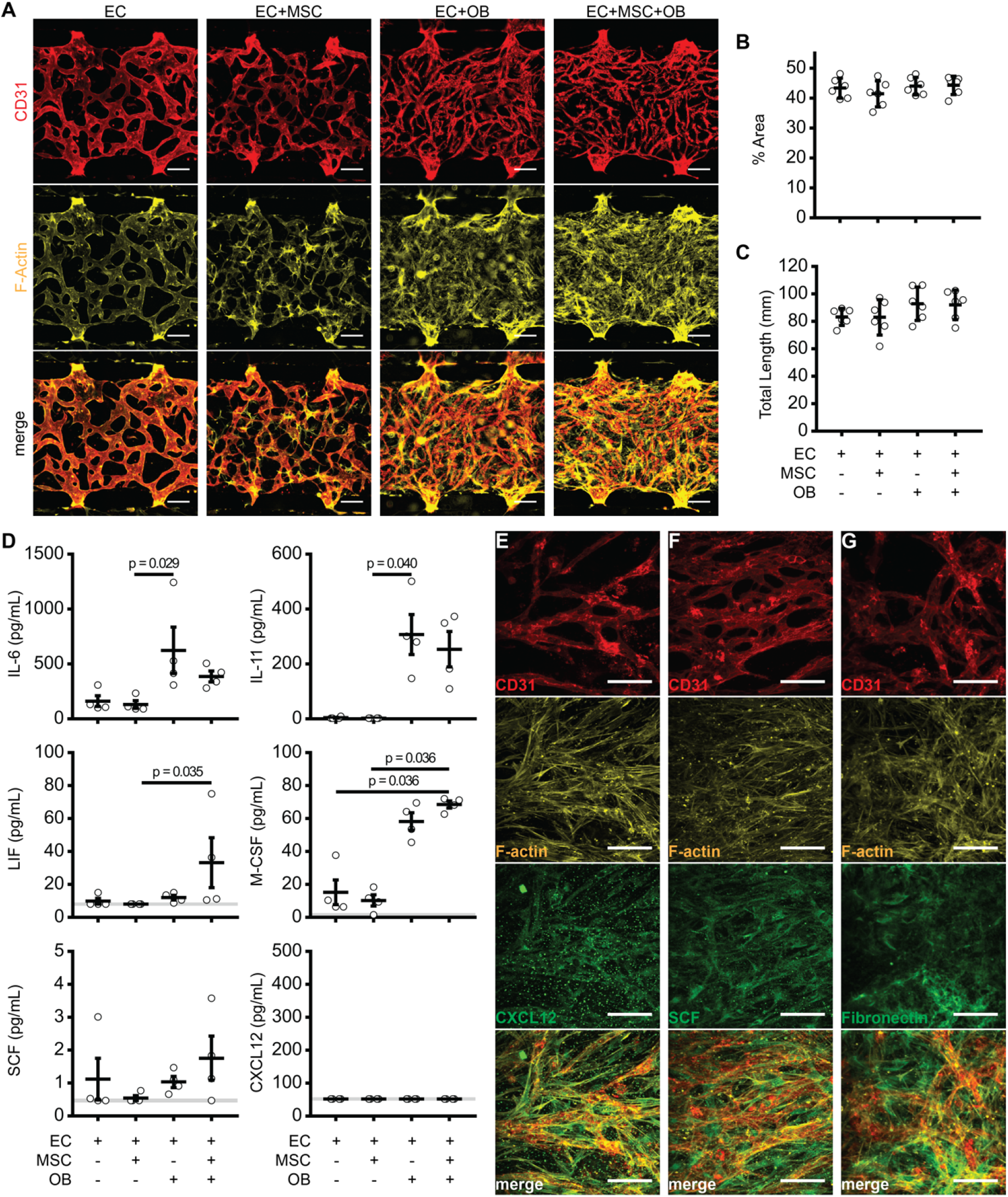
Vasculogenesis and cytokine expression in hBM-on-a-chip. hBM-on-a-chip with or without MSCs and OBs created using a fibrin (4 mg/mL) and collagen I (1 mg/mL) co-gel with VEGF (50 ng/mL, days 1-5) and Ang-1 (100 ng/mL, days 3-5) supplementation. (A) Immunofluorescence staining of CD31 (red) and F-actin (yellow). Scale bars: 100 μm. Quantification of the (B) percent area and (C) total length of vascular networks. Data are plotted as mean ± SD (n = 4-6 devices). Data were analyzed using a one-way ANOVA with Tukey’s post hoc test. No significance found at p < 0.05. (D) Hematopoietic cytokine secretion measured in device supernatant collected on day 5. Data are plotted as mean ± SEM (n = 4 devices). Data were analyzed using Kruskal-Wallis w/ Dunn’s post hoc test. Significance between groups (p < 0.05) is indicated in the figure. Immunofluorescence staining of (E) CXCL12, (F), SCF, (G) fibronectin after 5 days vasculogenesis in EC+MSC+OB hBM-on-a-chip. Scale bars: 100 μm.

### Effect of stromal cells on cytokine secretion

We observed differences in cytokine expression as function of stromal cell inclusion using multiplex cytokine detection to analyze the supernatant collected from the devices on day 5 of vasculogenesis (Fig. 3D). OB containing devices, in general, secreted higher amounts of cytokines. IL-6, a cytokine involved in B cell differentiation, was highly expressed in EC+OB (624±212 pg/mL) and EC+MSC+OB (386±49 pg/mL) samples, less was measured to be present in EC (162±49 pg/mL) and EC+MSC (132±34 pg/mL). IL-11, which is responsible for signaling during megakaryocyte maturation, was measure in trace concentrations in EC (4.1±1.8 pg/mL) devices and not at all in EC+MSC samples, while substantial concentrations were observed in EC+OB (308±73 pg/mL) and EC+MSC+OB (254±64 pg/mL) devices. Similarly, M-CSF, which induces macrophage differentiation of HSCs, was elevated in EC+OB (58.3±5.3 pg/mL) and EC+MSC+OB (68.6±2.1 pg/mL) devices, while little was detected in EC (15.2±7.6 pg/mL) and EC+MSC (10.3±3.5 pg/mL) devices. Relatively small concentrations of IL-7 (lymphoid progenitors), IL-34 (monocytes and macrophages), GM-CSF (granulocytes and macrophages), FLT-3L (dendritic cells), and SCF (HSC maintenance) were measured and there was no significant difference across groups. For both CXCL12 (hematopoietic chemoattractant) and IL-3 (myeloid progenitors), no measurable analytes were detected.

### Characterization of cytokine and ECM expression on the hBM-on-a-chip

After 5 days of vasculogenesis, hBM-on-a-chip containing the endosteal layer and subsequently seeded hMSCs and HUVECs were fixed and stained to characterize the presence and localization of cytokines and ECM relevant to the BM niche (Fig. 3E-G). CXCL12 and SCF were both found to be expressed by perivascular and endothelial cells. Fibronectin was observed to be present in the “central marrow” space outside of the newly formed vasculature. This arrangement of cytokine expression is consistent with the distribution that has been seen *in vivo*.

### HSPCs in hBM-on-a-chip

We next sought to investigate the inclusion of BM HSPCs in the hBM-on-a-chip and how MSCs and OBs affected their fate in our BM model. BM CD34+ cells were briefly expanded *in vitro* out of cryo-storage and then loaded into the central channel with HUVECs and MSCs. We measured the expansion of the HSPCs via microscopy over the 5 days of culture during vasculogenesis (Fig. 4A and Fig. S4A). On day 1, the number of HSPCs in hBM-on-a-chip was not significantly different in MSC and/or OB containing devices. By day 3 and continuing to day 5, devices without OBs had significantly more HSPCs than devices with the endosteal layer. HSPCs culture with ECs and MSCs expanded 2.41- and 2.28-fold, respectively, over 5 days, whereas both groups with HSPCs cultured in the presence of the pre-formed endosteal layer only expanded 1.83-fold (Fig. 4Aii).

**Fig. 4.**
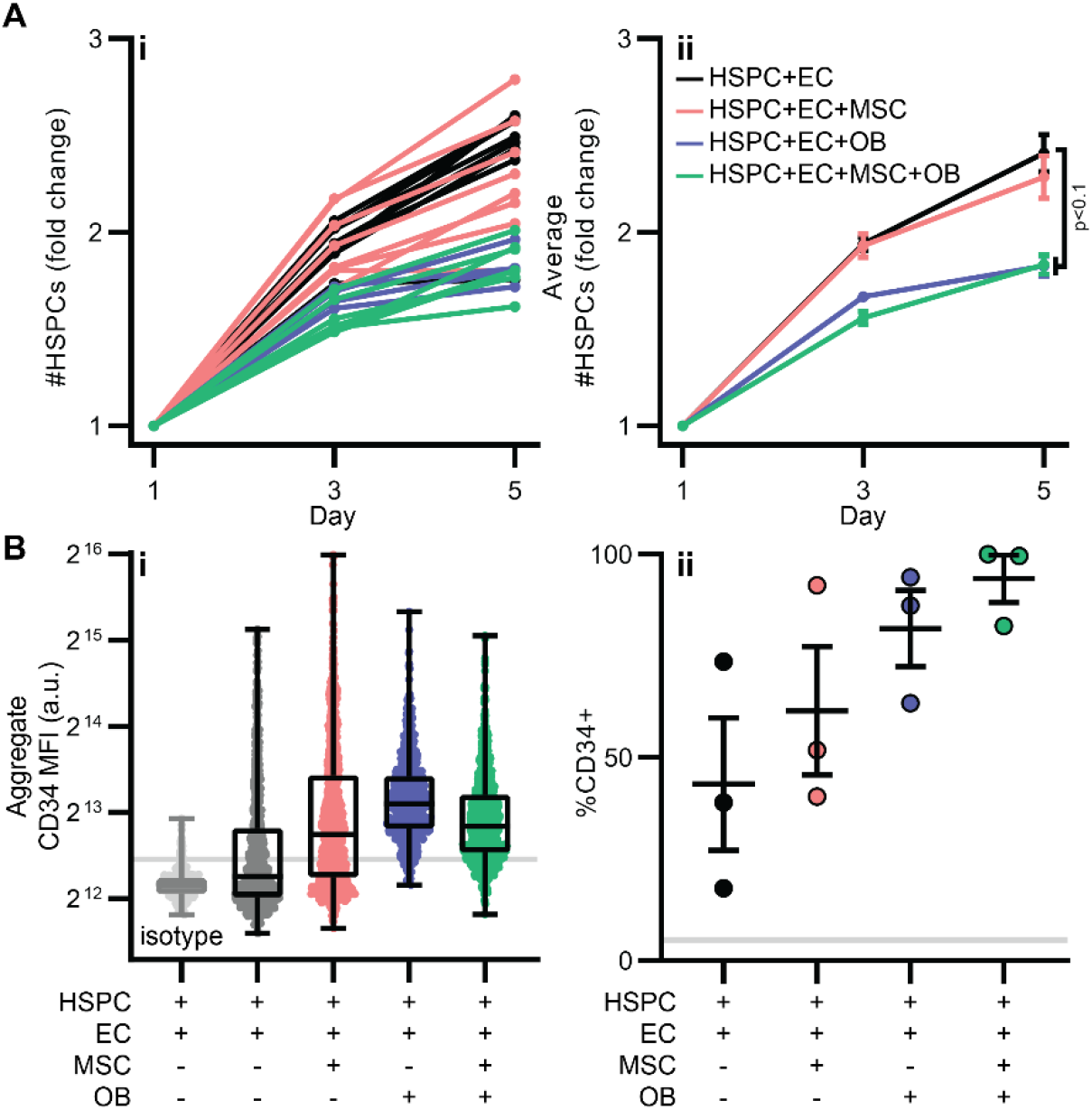
(A) Fold change number of HSPCs ((i) individual devices and (ii) summary data) on days 1, 3, and 5. Data are shown as mean ± SEM. (n = 4 devices EC+OB, n = 7 devices EC+MSC +OB, n = 8 devices EC and EC+MSC). Data were analyzed using Kruskal-Wallis w/ Dunn’s multiple comparisons test. EC vs EC+OB p = 0.0813; EC vs EC+MSC+OB p = 0.0162. (B) Quantification of immunofluorescence of HSPCs cultured for 5 days in hBM-on-a-chip with ECs only (black), with MSCs (red), OBs (blue), and both (green). (i) Aggregate MFI of CD34. Data are shown with median, quartiles, min and max (n = 452 cells for isotype from 1 device, n = 800 cells for EC, n = 907 cells EC+MSC, n = 945 cells EC+OB, n = 799 cells EC+MSC+OB pooled from 3 devices). (ii) Percentage of CD34^+^ cells. Data are shown as mean ± SEM (n = 3 devices). Grey line is percentage of CD34^+^ cells in isotype sample using gating scheme.

HSPCs will rapidly differentiate when cultured *in vitro*. We attempted to measure the effect of MSCs and OBs on the differentiation of HSPCs in the hBM-on-a-chip by staining for CD34 (Fig. 4B). HSPCs were pre-labeled with the membrane dye PKH67 to identify HSPCs for subsequent fluorescence quantification (Fig. S4B). We observed increased expression of CD34 in the presence of MSCs and/or OBs, as well as an increase in the percentage of CD34+ HSPCs.

### Mobilization of CD34+ HSPCs in hBM-on-a-chip

In order to measure the mobilization of HSPCs in hBM-on-a-chip, we designed an experimental assay where devices were imaged every 2 hours over a 24-hour period after treatment with mobilizing agents (Fig. S5). During each imaging session, devices were imaged 4 times in 5-minute intervals and these images were used to track cell movement and measure the length and displacement at discrete points in time during the 24-hour period. Untreated samples showed relatively steady movement of HSPCs when measured by either the length (Fig. S5B) or displacement (Fig. S5C) over 24 hours.

Treatment of hBM-on-a-chip (EC+MSC+OB+HSPC) with 10 ng/mL or 100 ng/mL G-CSF resulted in a moderate increase in the relative length and displacement of HSPCs at the lower concentration (Fig. 5A). After treatment with 10 ng/mL G-CSF, HSPCs had an increased tracked length between approximately 2-14 hours post-treatment. Untreated and samples mobilized with 100 ng/mL G-CSF did not show any sustained increase in HSPC tracked length but instead had a steady decrease from the peak at the 0-hour baseline measurement.

**Fig. 5.**
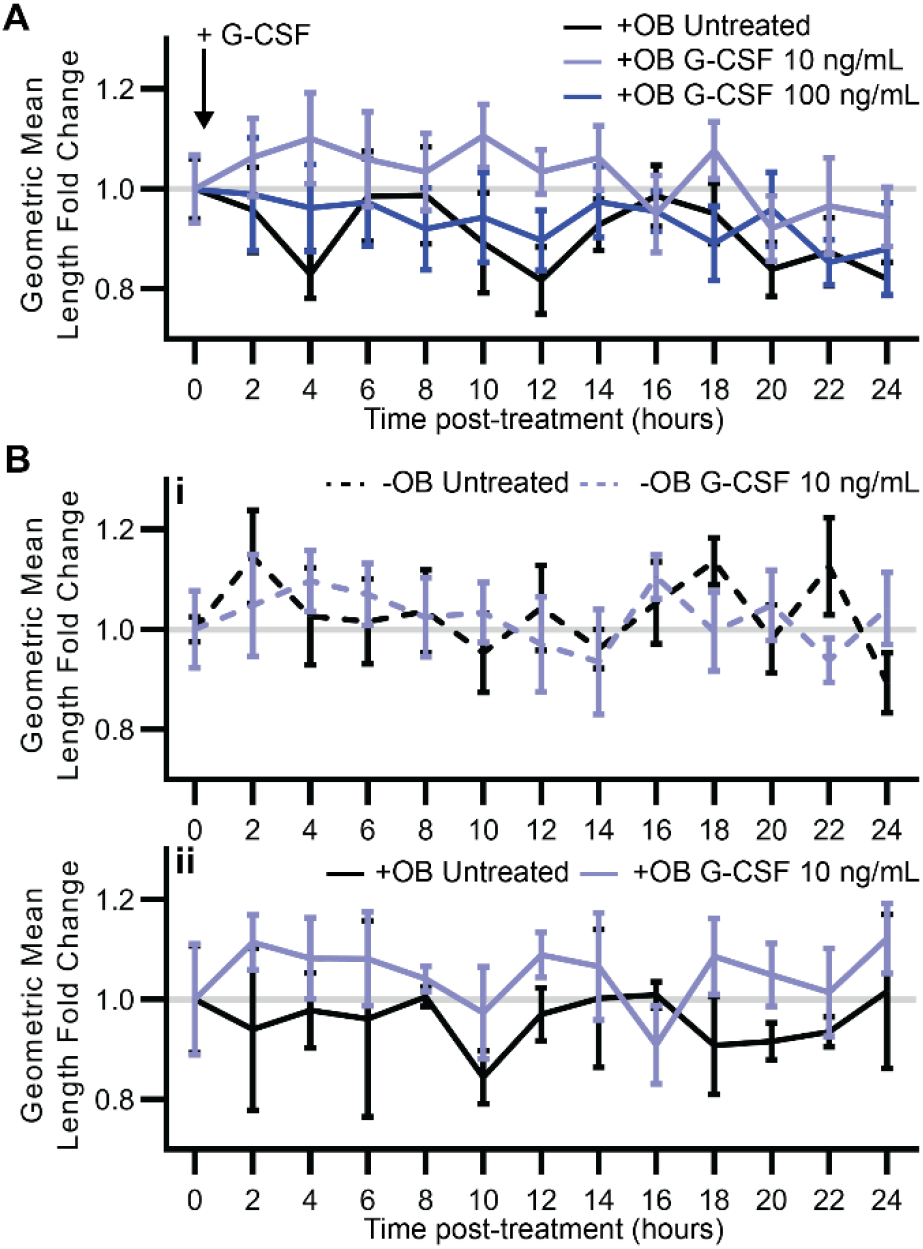
Mobilization of CD34+ BM HSPCs using G-CSF. hBM-on-a-chip (HSPC+EC+MSC+OB) were untreated (black) or treated with 10 ng/mL G-CSF (light blue) or 100 ng/mL G-CSF (dark blue) and movement of HSPCs was measured over 24-hours. (A) Geometric mean tracked length fold change during 15-minute imaging sessions at 2-hour intervals. Data are shown as mean of geometric means ± SEM (n = 7 devices untreated, n = 8 devices 10 ng/mL and 100 ng/mL G-CSF). (B) hBM-on-a-chip (i) without OBs (dotted lines) and (ii) with OBs (solid lines) were treated with 10 ng/mL G-CSF (light blue) or were untreated (black) and movement of HSPCs was measured over 24-hours. Data are shown as mean of geometric means ± SEM (n = 3 devices +OB untreated, n = 5 devices −OB untreated, n = 6 devices −OB and +OB 10 ng/mL G-CSF).

To determine whether the endosteal niche affected mobilization by G-CSF we measured mobilization in hBM-on-a-chip with OBs (a vascularized endosteal niche model) and without OBs (a perivascular niche model). Similar to the previous, dosing experiment (Fig. 5A), with an endosteal niche present (+OB) devices treated with 10 ng/mL G-CSF had increased HSPC track length from approximately 2-14 hours post-treatment when compared to the untreated +OB control (Fig. 5Bii). Without an endosteal niche, there was no trend in increased track length or displacement when compared to the untreated −OB control (Fig. 5i).

### Radiation of CD34+ HSPCs in hBM-on-a-chip

To investigate the effect of ionizing radiation on the BM microenvironment in hBM-on-a-chip, after 5 days of vasculogenesis, devices containing OBs, MSCs, and ECs were exposed to 0 Gy, 2.5 Gy, 5 Gy, or 10 Gy X-ray irradiation. Device supernatant was collected at 0 hours (pre-treatment) and 24 hours radiation exposure. To measure cytotoxicity because of radiation, the change in lactate dehydrogenase (LDH) activity of the supernatant after irradiation was measured (Fig. S6C). Although we did not measure a significant increase in LDH activity, the LDH activity in devices exposed to 5 Gy and 10 Gy X-ray radiation marginally increased after exposure, while the activity in devices exposed to 0 Gy and 2.5 Gy radiation slightly decreased during the same period.

The change in cytokine secretion because of X-ray exposure was also measured. IL-6, IL-7, IL-11, M-CSF, and SCF were detected, however CXCL12, IL-3, IL-15, IL-34, GM-CSF, FLT3L, and LIF were not above the detection limit of the assay. While there was no significant difference between the groups, it is worth noting that there was a decrease in secretion of IL-6, IL-7, IL-11, M-CSF, and SCF for all irradiated groups (Fig. S6D).

We sought to determine whether the presence of the endosteal niche was ameliorating the effects of radiation for HSPCs. hBM-on-a-chip with (EC+MSC+OB+HSPC) and without (EC+MSC+HSPC) the endosteal niche, were exposed to 5 Gy X-ray radiation and again apoptosis was measured 24 hours after exposure via a TUNEL assay (Fig. 6). Similarly, high backgrounds of apoptotic HSPCs were observed, 17.5 ± 0.1% of HSPCs in untreated −OB samples and 16.3 ± 2.8% of HSPCs in untreated +OB samples staining positive for TUNEL (Fig. 6B). While there was not a statistically significant difference between any of the groups, we observed a moderate increase in TUNEL^+^ HSPCs in +OB devices exposed to 5 Gy radiation (17.7 ± 2.0%). There was a larger increase in TUNEL^+^ HSPCs when exposed to 5 Gy radiation without the endosteal niche (-OB) present, with 22.6 ± 1.9% of HSPCs apoptotic.

**Fig. 6.**
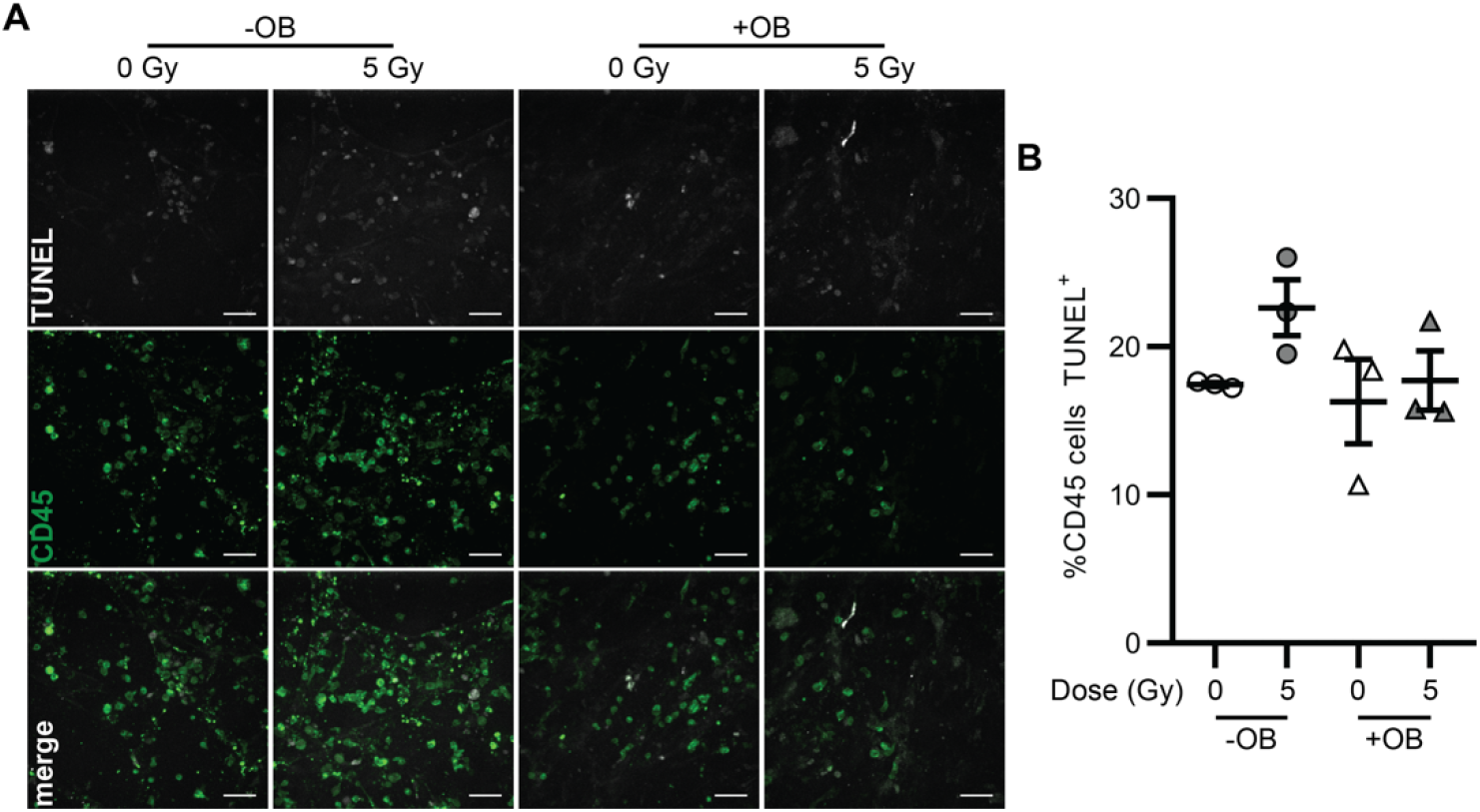
Effect of endosteal niche on ionizing radiation damage to HSPCs. (A) Representative images of TUNEL (white) and CD45 (green) staining of devices, with or without OBs, exposed to 0 or 5 Gy radiation 24 hours after exposure. Scale bar: 50 μm. (B) Quantification of percentage of TUNEL^+^ CD45 cells, with (triangles) or without (circles) OBs, 24 hours after 0 (white) or 5 (grey) Gy. Data are shown as mean ± SEM (n = 3 devices). Data analyzed using Kruskal-Wallis with Dunn’s multiple comparisons test. No significance between groups (p < 0.1).

## Discussion

We have developed a simple model of the hBM microenvironment that incorporates both the endosteal and perivascular niches and demonstrated that this approach for an hBM-on-a-chip generates basic cytokine and ECM expression characteristic of BM that has been observed *in vivo*. Additionally, we have shown that the inclusion of the endosteal layer and perivascular hMSCs impacts the proliferation and fate of HSPCs co-cultured within the device. Due to the limited volume of the current design, this model is not an appropriate approach for the *ex vivo* expansion of hematopoietic cells for eventual transplantation, however we believe the hBM-on-a-chip could be a useful tool for studying HSPC or other therapeutic cell interactions with specific BM microenvironments and the constituent stromal and hematopoietic cells. Inclusion of additional cell types, such as macrophages, osteoclasts, adipocytes, could easily be achieved and introduce additional nuance and complexity to this model. These studies could expand beyond normal conditions and the hBM-on-a-chip could be used as a platform for studying the effects of cancer, radiation, or other perturbed states, and for the screening of therapeutics that target BM resident cells to elicit a therapeutic response, such as mobilization.

The mobilization of HSPCs from the BM niche into peripheral blood for collection by apheresis is a critical process for in both autologous and allogeneic HSC transplantation (*35*). G-CSF has long been used in conjunction chemotherapy (typically cyclophosphamide) to mobilize BM HSPCs, however it is not effective in all patients. Previous chemotherapy or radiotherapy regimens, age, and disease burden are potential clinical variables that may cause poor mobilization (*36–38*). Over the last 20 years, AMD3100 (Mozobil, Plerixafor) has been developed and is now the second FDA approved mobilizing drug and is used, although at a significant increase in cost, as a second line treatment for patients who have failed G-CSF plus chemotherapy mobilization (*39*).

To measure the mobilization of HSPCs in hBM-on-a-chip, we developed a protocol that allowed for the periodic measurement of cell movement over a 24-hour period. We did so because we did not anticipate HSPCs would be drawn out of the hBM-on-a-chip, because there was no supplemented chemokine or existing gradient within the device that would direct the HSPCs in a specific direction. Rather, we hypothesized that mobilization would disrupt local signaling that was restricting HSPC movement and would therefore lead to an increase in either the magnitude or displacement of migration during a given time period.

While the mechanism of mobilization by AMD3100 is specific, G-CSF mobilization likely occurs through multiple pathways. G-CSF stimulates macrophages, osteoblasts, and osteoclasts to upregulate proteases (MMP9, cathepsins, etc.) that subsequently degrade CXCL12 and surface bound VLA-4 and CXCR4 on HSPCs (*40–45*). This disrupts the CXCL12-CXCR4 signaling axis and leads to an increase of HSPCs in peripheral blood. Because G-CSF mobilization is indirect and is mediated by stromal cells, onset of mobilization by G-CSF is relatively slow, taking more than 24 hours to reach maximum mobilization (*46*). AMD3100 is an antagonist of CXCR4 and directly competes with CXCL12 for binding. Consequently, AMD3100 mobilization is faster than G-CSF, with HSPCs elevated in peripheral blood as early as 30 minutes post injection (*47*). When used in combination with G-CSF, AMD3100 mobilizes more CD34+ HSPCs and has more predictable kinetics than G-CSF alone (*48–51*). Currently preclinical mobilization studies are restricted to animal models (*52*) and a sophisticated *in vitro* platform could be beneficial to screen novel therapeutics on a large population of human samples or in a patient specific BM model to assess an individual’s mobilization potential.

Mobilizing the HSPCs with G-CSF, we observed a dose specific response that lead to increased total length of migration and displacement over approximately a 14-hour period compared with the 0-hour baseline in both vascularized endosteal niche (+OB) and perivascular (-OB) hBM-on-a-chip devices. The mechanism of G-CSF mobilization is believed to go through any number of BM resident cells (neutrophils, osteoblasts, osteoclasts, macrophages) (*40–43*) with MSCs being a notable exception. The potential mobilization observed in samples without OBs is curious and, if true, suggests that either MSCs can mediate G-CSF induced disruption of CXCL12 signaling or that G-CSF is directly activating increased movement in HSPCs.

Ionizing radiation (IR) damages the BM microenvironment and resident HSPCs, and can occur both in accidental and clinical situations (*53*). IR harms HSPCs, results in the depletion of mature hematopoietic cells, and causes a range of symptoms associated with hematopoietic syndrome, and in extreme cases, fatality. While the impacts of IR on the hematopoietic system are well characterized (*54, 55*), the corresponding effects of IR on the HSPC niche and the BM microenvironment are less understood. MSCs are surprisingly somewhat resistant to the effects of IR and have been observed to potentially provide protection to other radiation damaged cells (*56*). Conversely, the activity of both osteoblasts and endothelial cells are altered by exposure to IR. Osteoblast activity is downregulated (*57–59*), decreasing the deposition of endosteal matrix and possibly, in conjunction with upregulation of osteoclasts, causing loss of bone mass at a larger scale (*60*). This could lead to the loss of the endosteal niche for hematopoietic progenitor cells that may be counteracted by increased osteogenic differentiation of MSCs (*61*). Common to radiation damage to all tissue types, the vasculature in BM is damaged and blood flow is disturbed (*57*). The perivascular niche for HSPCs is likely damaged and recovery of this microenvironment is essential for the regeneration of the hematopoietic system. The effects of radiation have, in the past, been studied mostly in animal models, recently organ-on-chip systems have been explored for their potential application in radiobiology (*62, 63*). Using organ-on-chip systems to examine the biologically response to radiation exposure not only moves preclinical studies away from animal models, but it may better recapitulate the response of human cells in radiation exposure situations.

Unexpectedly, exposure to relatively high doses of X-ray radiation (up to 10 Gy) did not induce a significant increase in cell death in vascularized endosteal niche hBM-on-a-chip (EC+MSC+OB), as measured by LDH release. There was a corresponding decrease, although not statistically significant, in cytokine expression 24 hours after radiation exposure. This suggested that the expression and, subsequently, the BM niche were potentially altered, albeit without widespread apoptosis of stromal cells. HSPCs cultured in the vascularized endosteal niche (HSPC+EC+MSC+OB) similarly did not exhibit a significant increase in apoptosis 24 hours after exposure as measured by damage to nuclear DNA (TUNEL). In comparison, we observed an increase, although not statistically significant, in TUNEL^+^ HSPCs in the perivascular niche (HSPC+EC+MSC) after 5 Gy radiation exposure. MSCs have been reported to be resistant to damage associated with radiation exposure (*56*) and given the greater number of MSC derived cells in OB containing devices, the density of these radio-resistant (and potentially radio protective) cells could be mitigating damage to HSPCs. Further study measuring the production of reactive oxygen species (ROS) after radiation exposure of the perivascular and vascularized endosteal niches may help explain the difference in HSPC outcome in these microenvironments.

## Materials and Methods

### Device design and fabrication

Photomask (CAD/Art Services) was designed using AutoCAD software (Autodesk). An SU-8 master mold was fabricated using previously described soft lithography techniques.(*64*) Briefly, SU-8 2150 (Microchem) was spun to a thickness of ~120 μm on a silicon wafer (University Wafers) using a G3P8 Spin Coater (SCS). SU-8 was exposed with UV light through the photomask using an MJB4 mask aligner (Suss Microtec). Uncrosslinked SU-8 was removed with SU-8 developer (Microchem) and silicon wafers were treated by vapor phase deposition of trichloro (1H, 1H, 2H, 2H-perfluorooctyl)silane (Sigma-Aldrich) to increase surface hydrophobicity.

Polydimethylsiloxane (PDMS) (Dow Corning) was mixed 10:1 (elastomer base: curing agent) and cast on silicone master mold. PDMS was cured at 65 °C. The PDMS layer containing the features was then removed from the master mold, a 3D printed reservoir mold was aligned on top and additional PMDS (10:1) was poured on top of the device to form media reservoirs. After curing at 65 °C, loading ports were made using 1 mm biopsy punch (Integra Miltex). Devices were bonded to a thin film (~300 um) of PDMS (5:1) using a plasma cleaner (Harrick Plasma). Prior to use, devices were washed with 70% EtOH.

To promote cell adhesion, the central gel channel was coated with 0.01% dopamine HCl (Sigma-Aldrich) in TE Buffer [pH 8.5] for 1 hour at room temperature, washed with PBS, coated with 100 μg/mL rat tail collagen I (Corning) in PBS for 1 hour at room temperature, washed with PBS, and then dried overnight at 65 °C.(*32*) Devices were sterilized by UV exposure for at least 30 minutes prior to culture of cells.

Detailed methods can be found in Supplementary Methods.

### Cell culture

Bone marrow derived human mesenchymal stem cells (hMSCs) (RoosterBio) were initially expanded in hMSC High Performance Media (RoosterBio) for a single passage. For subsequent passages, hMSCs were expanded in αMEM (Sigma-Aldrich) supplemented with 10% FBS (Hyclone) and 1% penicillin-streptomycin (Hyclone). hMSCs were used for culture in devices up to passage 6. Human umbilical vein endothelial cells (HUVECs) (Lonza) were expanded in EBM-2MV (Lonza) on tissue culture flasks coated with 0.1% gelatin (Sigma-Aldrich) and used up to passage 8. Human BM CD34+ cells (Lonza) were expanded for 5 days in Stemline II (Sigma-Aldrich) supplemented with 100 ng/mL SCF, TPO, and G-CSF (Peprotech). All cells were cultured at 37 °C and 5% CO_2_.

### Osteogenesis in hBM-on-a-chip

For the formation of the endosteal niche, hMSCs were seeded within the central gel channels of devices at a density of 5×10^5^ cells/mL. Cells were cultured within the devices for 21 days in αMEM osteogenic media (10% FBS, 1% penicillin-streptomycin, 10 mM β-glycerophosphate (Sigma-Aldrich), 50 μM ascorbic acid (Sigma-Aldrich), and 100 nM dexamethasone (Sigma-Aldrich)) with daily media exchange.

### Alizarin red and von Kossa staining

Osteogenic devices were washed with PBS, fixed with 4% formaldehyde in PBS for 15 minutes, and then washed with PBS. For Alizarin red staining, devices were washed twice with DI H_2_O and then stained for 5 minutes with 2% alizarin red (Sigma-Aldrich) in DI H_2_O [pH 4.1-4.3]. Alizarin red stain was removed by several washes with DI H_2_O, until liquid was clear. For von Kossa staining, devices were washed twice with DI H_2_O and then stained with 1% silver nitrate (Acros Organics) in DI H_2_O under a UV lamp for 15 minutes. Devices were washed twice with DI H_2_O and then incubated in 5% sodium thiosulfate (Acros Organics) in DI H_2_O for 5 minutes. Devices were then washed with DI H_2_O until liquid was clear. Stained devices were imaged using a Lionheart FX (BioTek Instruments). Color brightfield images were analyzed using open-source software ImageJ (https://imagej.nih.gov/ij/index.html). The red and green channels were used for von Kossa and Alizarin, respectively, to measure mean intensity and percent area.

### Vasculogenesis in hBM-on-a-chip

Vasculogenesis in central gel channel was accomplished using previously reported approaches (*33, 65*). Briefly, HUVECs (12×10^6^ cells/mL) and MSCs (6×10^5^ cells/mL) were suspended in EBM-2MV supplemented with thrombin (4 U/mL) (Sigma-Aldrich). A solution of fibrinogen (8 mg/mL) (Sigma-Aldrich) and collagen I (2 mg/mL) (Corning) in PBS was loaded into a central gel channel reservoir. The HUVEC/MSC cell suspension was added to fibrinogen solution (1:1) and mixed thoroughly. (Final cell suspension: HUVECs (6×10^6^ cells/mL), hMSCs (3×10^5^ cells/mL), thrombin (2 U/mL), fibrinogen (4 mg/mL), collagen I (1 mg/mL)). Immediately, the hMSC/HUVEC cell suspension was withdrawn from the opposite central gel port, drawing the cell suspension through the central gel channel. Devices were then incubated for 15 minutes at 37 °C, 5% CO_2_ to allow the fibrin gel to form. EBM-2MV was added to a reservoir on each side of gel channel and pulled through media channel, into connecting reservoir by using a micropipette to create negative pressure in the connecting 1-mm port. Cells were cultured with daily media exchange for 5 days to allow for vasculogenesis. Media was supplemented with VEGF (50 ng/mL) on day 2 and with VEGF (50 ng/mL) and angiopoietin-1 (ANG-1) (100 ng/mL) (Peprotech) on days 3, 4, and 5.

### Immunofluorescence staining

Staining procedure for devices was adapted from Chen *et al* 2007 (*27*). Devices were washed with PBS, fixed with 4% formaldehyde (ThermoFisher), and permeabilized with 0.1% Triton X-100. Prior to staining, cells were blocked with 5% BSA, 3% goat serum in PBS. Primary antibodies were diluted (1:100) in blocking buffer and devices were stained overnight at 4 °C. Devices were then washed with 0.1% BSA in PBS and stained with secondary antibodies (1:200) and Phalloidin AF647 (1:40) diluted in wash buffer for 3 hours at RT. Devices were washed with wash buffer and stored at 4 °C until imaging. For immunofluorescence imaging of cytokines and ECM in full hBM-on-a-chip devices, samples were imaged using an UltraVIEW VoX spinning disk confocal microscope (PerkinElmer). For detailed information on antibodies, see Table S2.

### Vascular network analysis

Devices stained with Alexa Fluor 647 anti-human CD31 (BioLegend) and Alexa Fluor 594 Phalloidin (ThermoFisher) were imaged using a Lionheart FX (BioTek Instruments). Images were processed using open-source software ImageJ and contrast corrected images were analyzed using Angiotool(*66*) to measure percent area and total network length.

### Multiplex cytokine detection

Media was collected from devices at designated time by collecting media from one side of device, waiting for 5 minutes to allow for gravity driven flow through the device and then collection of all media from reservoirs. Device media was immediately stored on ice and then flash frozen in liquid N_2_ for storage prior to analysis. Samples were thawed on ice prior to detection and analysis using LEGENDplex human hematopoietic stem cell panel (BioLegend), according to manufacturer’s protocol.

### CD34+ HSPCs culture in hBM-on-a-chip

BM CD34+ HSPCs were labelled with PKH67 green fluorescent cell stain (Sigma-Aldrich) and loaded at a final concentration of 3×10^5^ (20:1 HUVEC:HSPC ratio, 6×10^5^ HSPCs/mL in thrombin cell suspension prior to mixing with fibrinogen). This concentration results in ~500 HSPCs within the central channel. Devices were imaged on days 1, 3, and 5 after loading using a Lionheart FX (BioTek Instruments). Images were analyzed using Gen5 (BioTek Instruments) to count the number of HSPCs and progenitors at each time point.

### Mobilization of HSPCs

After 5 days of culture, hBM-on-a-chips were imaged periodically to measure the “mobilization” of CD34+ HSPCs. Baseline measurements were made at 0 hours, after which supernatants were collected and was replaced with media supplemented with mobilizing agents at indicated concentrations. Samples were imaged at intervals of 2 hours for 24 hours for a total of 13 image sessions. During each image session, devices were imaged at intervals of 5 minutes for 15 minutes, for a total of 4 time points. Samples were imaged using a Cytation 3 (BioTek Instruments) and automated imaging of multiple plates was achieved using a BioSpa 3 (BioTek Instruments). Image sequences were imported into Volocity software (PerkinElmer), cells were tracked within each image session, and the length and displacement of individual cells were measured at each time point.

### Radiation exposure

On day 5, hBM-on-a-chip devices were transported to an animal facility and exposed to ionizing radiation using an RS 2000 X-ray Irradiator (Rad Source). To control for effects of transportation, untreated samples were also transported to the facility. The duration of the process (transportation to and from the facility and X-ray exposure) was ~1 hour, during which time the samples were not held at 37 °C, 5% CO_2_.

### TUNEL assay

DNA damage to HSPCs was observed using terminal deoxynucleotidyl transferase-mediated dUTP-biotin nick-end labelling (TUNEL) immunostaining (Click-iT TUNEL Alexa Fluor Assay Kit, ThermoFisher) according to the manufacturer’s protocol. Briefly, formaldehyde fixed devices were first permeabilized with Triton-X100 (Avocado). Next, terminal deoxynucleotidyl transferase (TdT) was used to incorporate dUTP into double stranded breaks of DNA. Alexa Fluor 647 azide was then bound to dUTP using click chemistry. As membrane stains of HSPCs do not survive Triton-X100 permeabilization, HSPCs were then stained overnight using anti-human CD45 Alexa Fluor 488 (BioLegend). Samples were imaged using a spinning disk confocal microscope (PerkinElmer) and images were analyzed using Volocity software (PerkinElmer).

### Statistical analysis

Sample sizes are indicated in the figure captions and individual data points are shown. GraphPad Prism was used for statistical analysis. Groups were tested for normality using the Shapiro-Wilk normality test. If all groups in an experiment passed normality test, one-way ANOVA with Tukey’s multiple comparison test was used. If a single group did not pass normality test, Kruskal-Wallis with Dunn’s multiple comparison test was used.

## H2: Supplementary Materials

Supplementary Methods

Fig. S1. Detailed schematic of hBM-on-a-chip design and fabrication.

Fig. S2. ECM coating to promote cellular adhesion to PDMS and “survival” of device air-liquid interface during culture.

Fig. S3. Optimization of cytokine and hydrogel conditions for vasculogenesis.

Fig. S4. Effect of stromal cells on CD34+ BM HSPCs in hBM-on-a-chip.

Fig. S5. Measuring mobilization in hBM-on-a-chip.

Fig. S6. Effective X-ray radiation dose and effect of radiation on cytokine secretion in hBM-on-a-chip.

Table S1. List of materials.

Table S2. Antibodies and dilutions.

Table S3. Primary cell sources.

## Acknowledgements

We would like to acknowledge the Petit Institute Core Facilities for their services and shared resources that enabled us to produce this publication.

## Funding

This research was supported by funding from the Georgia Tech Foundation (to K.R.), Georgia Tech Research Alliance (to K.R.), the Marcus foundation (to K.R.), NSF Engineering Research Center Grant (NSF EEC 1648035 to K.R.), and NSF Graduate Research Fellowship under Grant No. DGE-1148903 (to M.R.N., D.G. and J.C.M.).

## Author contributions

M.R.N. designed and performed experiments, analyzed data, wrote the manuscript. D.G, J.C.M., D.F.R., and E.K. performed experiments, analyzed data, and edited the manuscript. KR designed experiments and edited the manuscript.

## Competing interests

The authors declare that they have no competing interests.

## Data and materials availability

All data needed to evaluate the conclusions in the paper are present in the paper and/or in the Supplementary Materials. Additional data related to this paper may be requested from the authors. Correspondence and requests for materials should be addressed to K.R.

## Notes

#### Summary of Updates

Figure 1

