## Supplemental Methods and Figures for "A Multi-Niche Microvascularized Human Bone-Marrow-on-a-Chip"

### Supplementary Materials

### Supplementary Methods

**Burst pressure calculations**

Burst pressure was calculated using an approached described by Wang *et al*. 2016, which is briefly described below (*67*). The pressure difference at the air-liquid interface for advancing liquid within the gel channel can be represented by the Young-Laplace equation:

$$P_{liquid\ advance} - P_{air} = -2\gamma\left(\frac{cos\theta_A}{w_{channel}} + \frac{cos\theta_A}{h}\right)$$

Where $P_{liquid\ advance}$ is the liquid pressure inside the gel channel, $\gamma$ is surface tension ($\gamma$ = 0.072 N m$^{-1}$), $\theta_A$ is the critical contact angle where liquid will burst or advance ($\theta_A$ = 140º), $w$ is the width of the channel, and $h$ is the height of the channel. The advancing pressure for liquid within gel channel ($w_{channel}$ = 1000 µm, $h$ = 150 µm) was found to be 845 Pa or 86 $mmH_2O$. The burst pressure at the air-liquid interface at the pores is represented as:

$$P_{liquid\ burst} - P_{air} = -2\gamma\left(\frac{cos\theta_A^*}{w_{pore}} + \frac{cos\theta_A}{h}\right)$$

Where $\theta_A^*$ is the contact angle of the liquid with the inner facing side wall of the channel dividers (however, $\theta_A^*$ is limited to 180º, the maximum contact angle for a liquid meniscus). The burst pressure at the communication pores ($w_{pore}$ = 50 µm, $h$ = 150 µm) is 3615 Pa or 369 $mmH_2O$. The difference between $P_{liquid\ advance}$ and $P_{liquid\ burst}$ is 2770 Pa or 282 $mmH_2O$.

**Detailed v1.0 device fabrication**

PDMS (Dow Corning) was mixed 10:1 (elastomer base: curing agent) and cast on SU-8 master mold. PDMS was cured at 65 °C. The PDMS layer containing the features was then removed from the master mold and loading ports were made using a 1 mm biopsy punch (Integra Miltex). To form the media reservoir layer, PDMS was mixed 10:1 and cast in a 100 mm petri dish to a thickness of 5-6 mm. PDMS was cured at 65 °C. The thick disc of PDMS was then cut into single-device sized squares and media reservoirs were made using a 4 mm biopsy punch. The top surface of the device layer was then bonded to the media reservoir layer using a plasma cleaner (Harrick Plasma). To form the PDMS coated coverslips, PDMS was mixed 10:1 and 50 µL were applied to a clean glass coverslip, the coverslip was then sheared against a glass slide to evenly coat the surface and the PDMS was cured at 65 °C. The PDMS coated coverslip was then bonded to the bottom surface of the device layer using a plasma cleaner (Fig. S1D,E). Prior to use, devices were washed with 70% EtOH.

**Detailed v1.1 device fabrication**

PDMS was mixed 10:1 and cast on SU-8 master mold. PDMS was cured at 65 ºC. The PDMS layer containing the features was then removed from the master mold, a 3D printed reservoir mold was aligned on top and additional PMDS (10:1) was poured on top of the device to form media reservoirs. After curing at 65 ºC, loading ports were made using 1 mm biopsy punch. A thin film of PDMS was made by mixing PDMS 5:1, casting a thin layer (~300 µm) in a 150 mm petri dish and curing at 65 °C. Devices were bonded to the thin film of PDMS using a plasma cleaner, and individual devices were cut for use in cell culture (Fig. S1F, G). Prior to use, devices were washed with 70% EtOH.

**Detailed v2.0 device fabrication**

hBM-on-a-chip was integrated into a standard well-plate format using previously published methods (Fig. S1H, I) (*29-31*). Because a standard 100 mm silicon wafer does not have enough area to pattern the entire 4x2 hBM-on-a-chip array (Fig. S1J, K), a polyurethane master mold was first fabricated from PDMS cast on a SU-8 master. PDMS was mixed 10:1 (elastomer base: curing agent) and cast on the SU-8 master mold () to create two copies of the 2x2 array. The two pieces were aligned, feature-side down, and made into a single block by casting in additional PDMS. A polyurethane master mold was then cast on the single PDMS piece containing the 4x2 array using a 2-part polyurethane liquid plastic (Smooth Cast 310, Smooth-On Inc.) (*30*). To prevent adhesion of PDMS subsequently cast, the polyurethane master mold was treated by vapor phase deposition of trichloro (1H, 1H, 2H, 2H-perfluorooctyl) silane (Sigma-Aldrich).

PDMS was mixed 10:1, cast on the polyurethane master mold, and cured at 65 °C. The PDMS layer containing the features was then removed from the master mold and loading ports were made using a 1 mm biopsy punch. A thin film of PDMS was made by mixing PDMS 5:1, casting a thin layer (~600 µm) in a 150 mm petri dish and curing at 65 °C. The feature layer of PDMS was then bonded to the PDMS film using a plasma cleaner. The bonded PDMS devices were then attached to a bottomless 96-well plate (Greiner Bio-One) using a chemical gluing method (*31*). Briefly, the 96-well plate was immersed in 2% (v/v) 3-mercaptopropyl trimethoxysilane (Sigma-Aldrich) in methanol for 1 minute, rinsed with deionized $H_2O$, and dried. The bonded PDMS devices were then plasma bonded to the 96-well plate using a plasma cleaner. To provide support to the bottom PDMS surface, glass coverslips (Fisher Scientific) were adhered to the PDMS film by plasma bonding. Prior to use, devices were washed with 70% EtOH and DI $H_2O$.

**PDMS ECM Coating**

**Vascular perfusion**

To visualize perfusion of microvasculature, 70 kDa dextran-FITC (Sigma-Aldrich) was flowed through the media channel on one side of the device. After 5 days of vasculogenesis, media was aspirated from all hBM-on-a-chip media ports (EC+MSC) and replaced with 50 µL of 10 µg/mL dextran-FITC in EGM-2MV in two, connected media ports. Devices were immediately imaged using a Lionheart FX (BioTek Instruments) microscope to visualize the flow of dextran through the central channel of the device.

**Radiation dose measurement and PDMS attenuation**

Dosimeters (nanoDot™, Landauer) were exposed to theoretical radiation doses of 2.5, 5, and 10 Gy using an RS 2000 X-ray Irradiator (Rad Source) at 2.15 Gy/min. To determine X-ray attenuation by the PDMS in hBM-on-a-chip, dosimeters were placed underneath a well-plate hBM-on-a-chip during exposure. Control samples were exposed without PDMS device shielding. A transit control dosimeter was also measured and the background radiation dose during shipping and use was subtracted from all test dosimeters. Dosimeters were returned to the service provider (Landauer) for dose measurements.

To determine the X-ray attenuation of hBM-on-a-chip, dosimeters were exposed to X-ray radiation doses of 2.5 Gy, 5 Gy, and 10 Gy through the well-plate hBM-on-a-chip device and compared to un-shielded, control dosimeters (Fig. S6A). Dosimeters shielded by the device did not show any reduction in dose compared to the control dosimeters. The measured dose of radiation was less than the theoretical dose for all three groups. The measured dose (both control and PDMS shielded) was 1.73 ± 0.04 Gy, 3.89 ± 0.03 Gy, and 8.03 ± 0.11 Gy (mean ± SEM) for the 2.5 Gy, 5 Gy, and 10 Gy groups, respectively. This corresponds to yields of 69%, 78%, and 80% of the theoretical doses.

Grouping the unshielded control samples and the PDMS shielded samples together (Fig. S6B), linear regression was performed to determine the difference between the theoretical radiation dose and the actual radiation dose measured:

$$Dose_{Measured} = 0.8108\, Dose_{Theoretical} - 0.1349\, ; R^2 = 0.9987$$

Exposure settings were subsequently adjusted to account for discrepancy between the irradiator's exposure rate and the measured doses.

**Lactate dehydrogenase (LDH) assay**

Device supernatant was collected (0 hour), replaced prior to radiation exposure and collected 24 hours after exposure. Supernatant was flash frozen immediately upon collection and stored at -20

°C prior to analysis. LDH activity was measured using the Pierce™ LDH Cytotoxicity Assay (ThermoFisher) according to the manufacturer's protocol. For each device, *i,* the 24-hour fold-change in LDH activity was calculated by dividing the background subtracted $A_{490}$ at 24 hours from the sample matched 0-hour value.

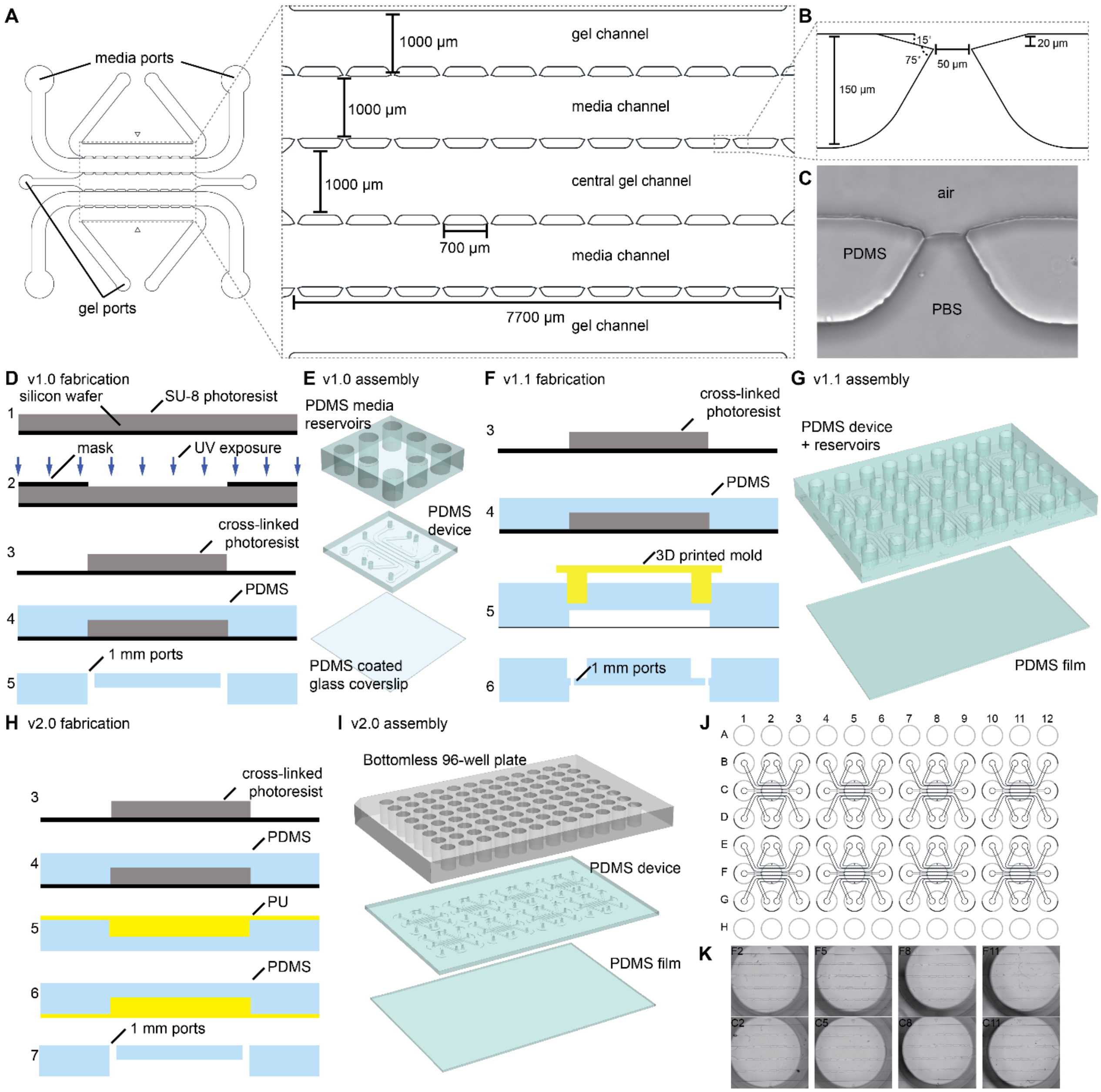


**Fig. S1. Detailed schematic of hBM-on-a-chip design and fabrication.** (A) The 5-channel PDMS microfluidic device was accessible through 1-mm gel loading ports (for central and outer channels) and media loading ports. Each channel has a width of 1000 μm and the channels are connected via 11 communication pores that are formed by 10 posts. (B) The communication pores are 50 μm in width and are designed to enable consistent formation of air-liquid-interface. (C) Example of air-liquid interface formed at a communication pore. (D) For v1.0, PDMS features were fabricated using standard soft lithography techniques. (1) A silicon wafer was spun coat with SU-8 photoresist, (2) exposed to UV light through a patterned photomask and (3) then the uncrosslinked SU-8 was removed leaving the patterned, SU-8 master mold. Then, (4) PDMS was cast on the SU-8 master mold and loading ports were created in the PDMS device layer. (E) The PDMS device layer was bonded to a PDMS media reservoir layer and to a PDMS coated coverslip to create a finished device. (F) For v1.1, the initial steps of fabrication (1-4) were unchanged from version 1.0. (5) To form the media reservoirs, a 3D printed mold was placed on top of the PDMS device layer and additional PDMS was cast to form the media reservoirs, after which (6) loading ports were made. (G) The PDMS device/media reservoir layer was plasma bonded to a thin PDMS film and cut into individual devices. (H) For v2.0 fabrication, the initial steps of fabrication (1-4) were unchanged from v1.0. To assemble a 2x4 array of devices, a polyurethane master mold was cast from two 2x2 device arrays. (I) The device layer of PDMS was first bonded to a thin PDMS film, then using a chemical gluing technique, the device layer was bonded to a bottomless 96-well plate. (J) The resulting device created a 2x4 array of devices that utilizes the wells of the well plate as media reservoirs, loading ports, and imaging windows. (K) Alignment of the devices creates 8 uniform devices in a single, standard well plate format.

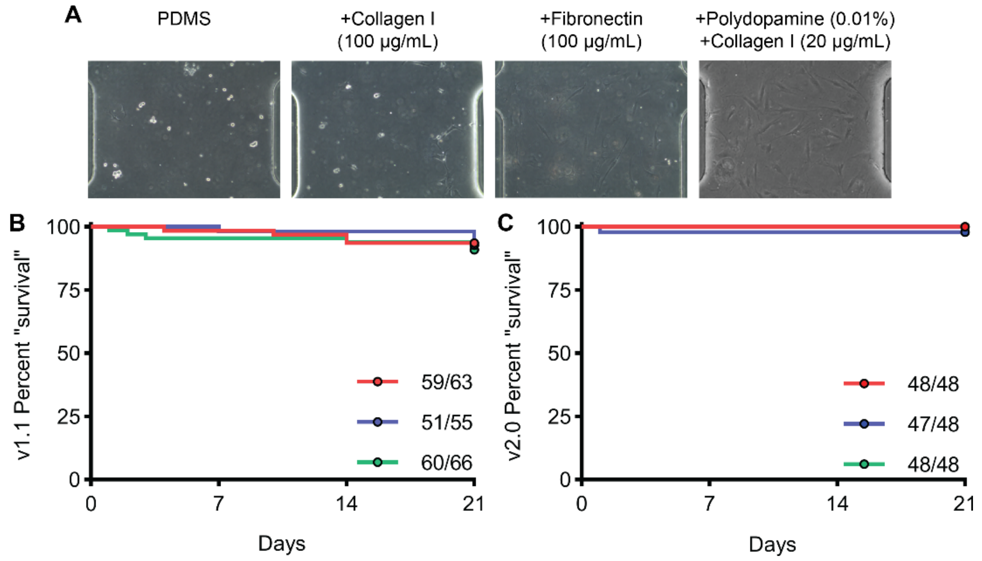


**Fig. S2. ECM coating to promote cellular adhesion to PDMS and "survival" of device air-liquid interface during culture.** (A) Phase contrast images of MSCs 24 hours after seeding in PDMS hBM-on-a-chip devices that were uncoated or coated with collagen 1 (100 µg/mL), fibronectin (100 µg/mL), or PDA (0.01%) and collagen I (20 µg/mL). "Survival" of viable devices using (B) v1.1 and (C) v2.0 hBM-on-a-chip device. Devices "fail" when media leaks from the central channel into adjacent media channels, rendering it impossible to isolate the central channel when loading cells on top of the endosteal layer.

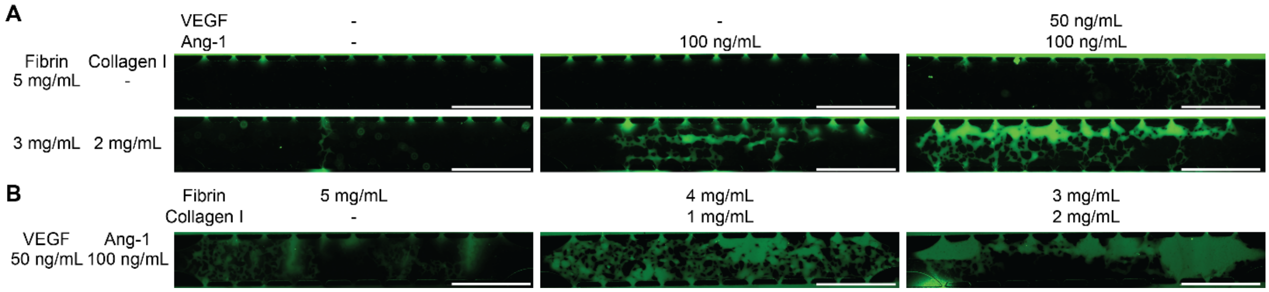


**Fig. S3. Optimization of cytokine and hydrogel conditions for vasculogenesis.**(A) Perfusion of vascular networks with 70 kDa dextran-FITC in hBM-on-a-chip (EC+MSC) using fibrin only (5 mg/mL) or fibrin-collagen co-gel (3 mg/mL, 2 mg/mL), with and without VEGF (50 ng/mL) and Ang-1 (100 ng/mL) supplementation. (B) Perfusion of vascular networks with 70 kDa dextran FITC in hBM-on-a-chip (without OB) with VEGF (50 ng/mL) and Ang-1 (100 ng/mL) supplementation, using fibrin only (5 mg/mL) or fibrin-collagen co-gels at 4:1 or 3:2 ratios Scale bar: 1000 μm.

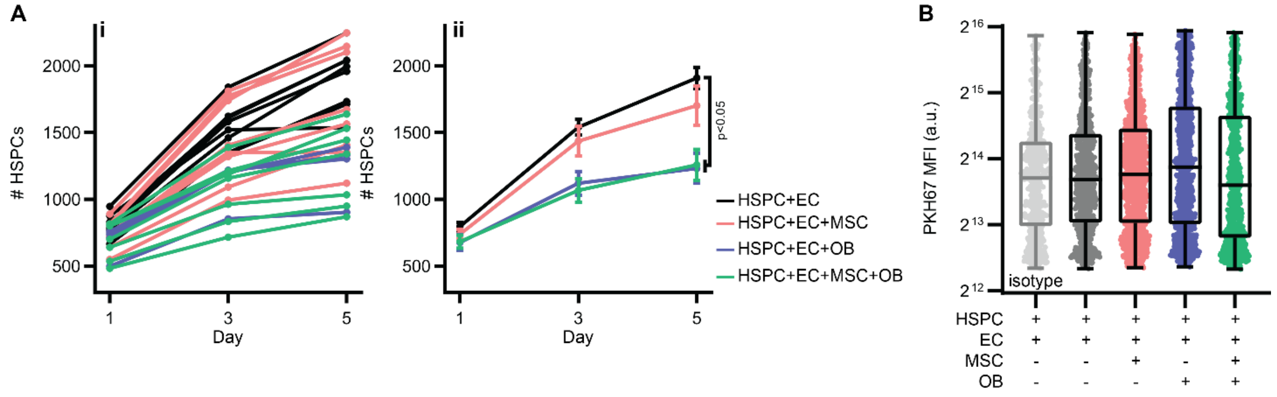


**Fig. S4. Effect of stromal cells on CD34$^+$ BM HSPCs in hBM-on-a-chip.** (A) Number of HSPCs ((i) individual devices and (ii) summary data) in hBM-on-a-chip with ECs only (black), with MSCs (red), OBs (blue), and both (green) on days 1, 3, and 5. Data were analyzed using Kruskal-Wallis w/ Dunn's multiple comparisons test. EC vs EC+OB p = 0.0210; EC vs EC+MSC+OB p = 0.0166. Quantification of immunofluorescence of HSPCs cultured for 5 days in hBM-on-a-chip with ECs only (black), with MSCs (red), OBs (blue), and both (green). (B) MFI of PK467 (cell tracker). Data are shown with median, quartiles, min and max (n = 452 cells for isotype from 1 device, n = 800 cells for EC, n = 907 cells EC+MSC, n = 945 cells EC+OB, n = 799 cells EC+MSC+OB pooled from 3 devices).

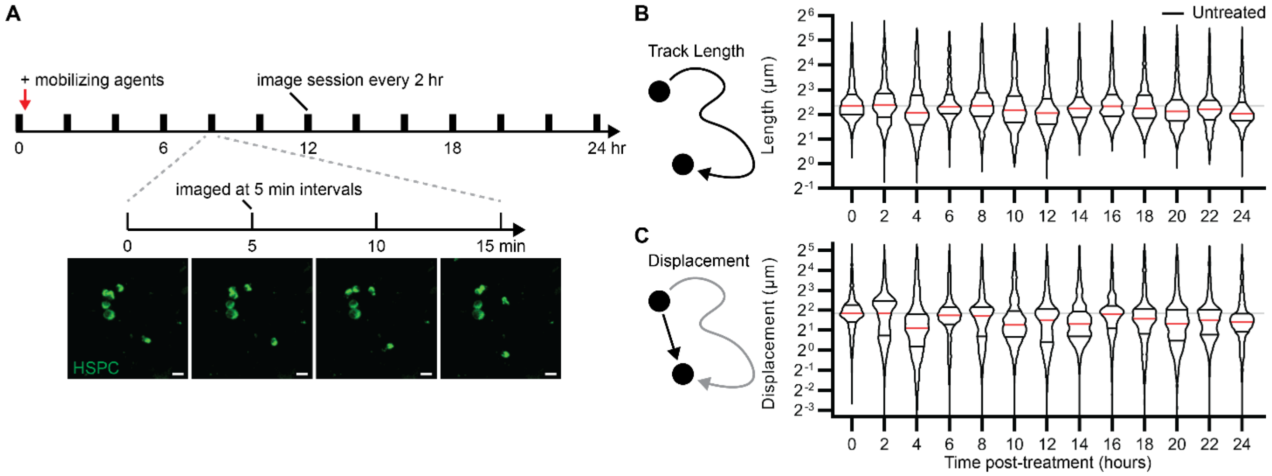


**Fig. S5. Measuring mobilization in hBM-on-a-chip.**(A) To observe the "mobilization" of HSPCs, we imaged devices 13 times over a 24-hour period. After the first imaging session, media was collected from the devices and it was exchanged for untreated media or media supplemented with mobilizing agents. During each imaging session, devices were imaged 4 times in 5-minute intervals. Cells were tracked during each imaging session to measure the track length and displacement at each time point. Scale bar: 20 µm. (B) Track length and (C) displacement over 24 hours in untreated devices. (B) and (C): data (n = 853-1236 tracked cells per time point pooled from 7 devices) are shown with median (red), quartiles (black), and 0-hour median (gray line).

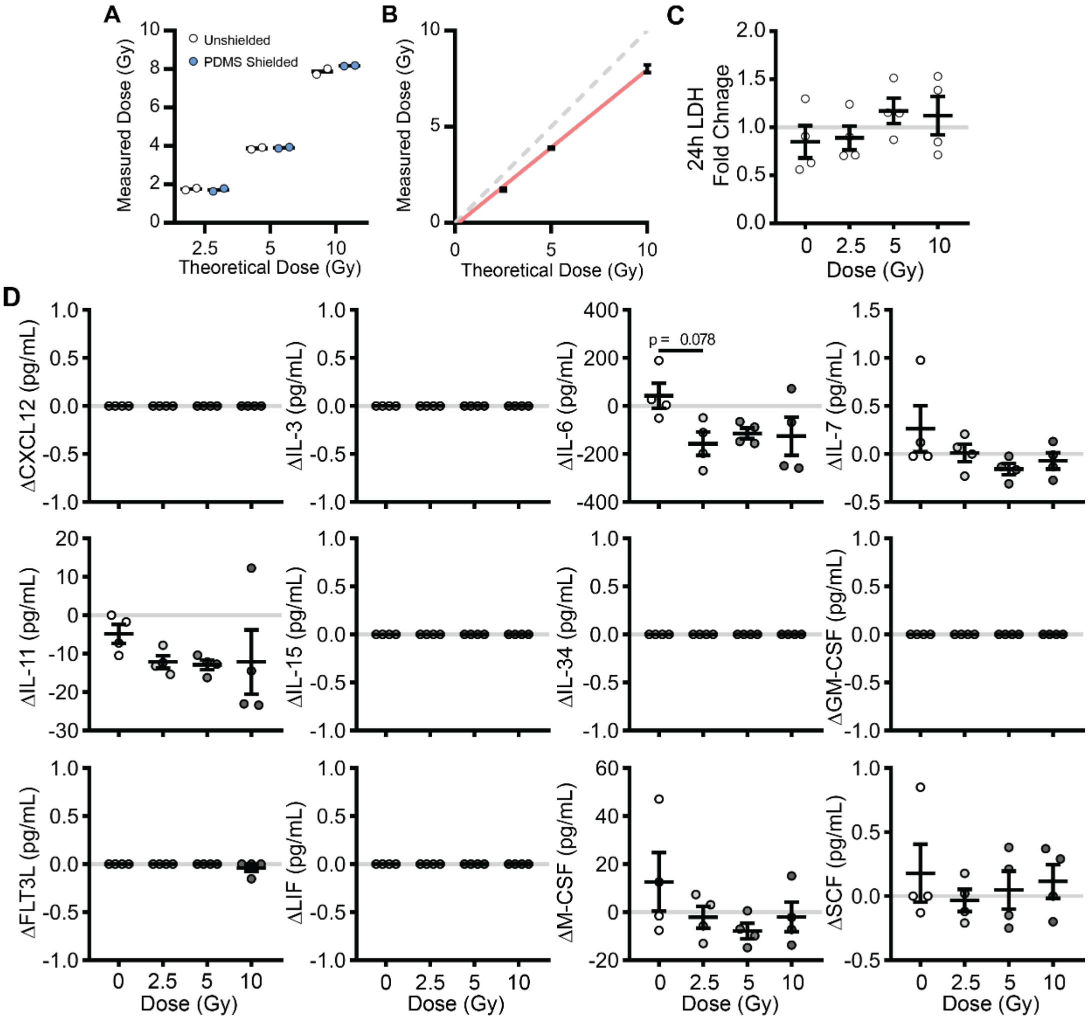


**Fig. S6. Effective X-ray radiation dose and effect of radiation on cytokine secretion in hBM-on-a-chip.**(A) Measured radiation dose of NanoDot® dosimeters exposed to 2.5, 5, or 10 Gy X-ray radiation with (blue) or without (white) PDMS shielding. Data is shown as mean (n = 2 dosimeters). (B) Linear regression of grouped (unshielded and PDMS shielded) dosimeter readings (red) used to interpolate exposure settings for accurate dosing. Data are shown as mean ± SD (n = 4 dosimeters). Perfect correlation (y = x) between theoretical and measured dose is shown by grey dotted line. Fold change in LDH released from hBM-on-a-chip (EC+MSC+OB) 24 hours post-X-ray irradiation compared to baseline (0 hours). Data are shown as mean ± SEM (n = 4 devices). Data analyzed using Kruskal-Wallis with Dunn's multiple comparison test. No significance between groups ($p < 0.1$). Change in hematopoietic cytokine measured in hBM-on-a-chip (EC+MSC+OB) supernatant before (0 hours) and 24 hours after devices were exposed to 0 (white), 2.5 (light grey), 5 (grey), or 10 (dark grey) Gy X-ray radiation. Data are shown as mean ± SEM (n = 4 devices). Data analyzed using Kruskal-Wallis with Dunn's multiple comparisons test.

**Table S1. List of materials.**

| Category | Name | Supplier | Catalog Number(s) |
|---|---|---|---|
| Chemical | β-Glycerophosphate | Sigma-Aldrich | G9422 |
| Chemical | (3-mercaptopropyl) trimethoxy silane | Sigma-Aldrich | 175617 |
| Chemical | Alizarin Red S | Sigma-Aldrich | A553325G |
| Chemical | AMD3100 | Sigma-Aldrich | A5602 |
| Chemical | Bovine gelatin | Sigma-Aldrich | G9391 |
| Chemical | Bovine serum albumin | Sigma-Aldrich | A2153 |
| Chemical | Dexamethasone | Sigma-Aldrich | D2915 |
| Chemical | Dopamine hydrochloride | Sigma-Aldrich | H8502 |
| Chemical | Goat serum | ThermoFisher | 10000C |
| Chemical | L-Ascorbic Acid | Sigma-Aldrich | 255564 |
| Chemical | PKH67 | Sigma-Aldrich | MIDI67 |
| Chemical | Rat Tail Collagen I | Corning | 354249 |
| Chemical | Silver Nitrate | Acros Organics | 197680250 |
| Chemical | Sodium Thiosulfate | Acros Organics | 202870010 |
| Chemical | Trichloro (1*H*,1*H*,2*H*,2*H*-perfluorooctyl) silane | Sigma-Aldrich | 448931 |
| Chemical | Triton X-100 | Avocado | A16046 |
| Cytokine | Human Ang-1 | PeproTech | 130-06 |
| Cytokine | Human CXCL12 | Abcam | ab9798 |
| Cytokine | Human G-CSF | PeproTech | 300-23 |
| Cytokine | Human SCF | PeproTech | 300-07 |
| Cytokine | Human TPO | PeproTech | 300-18 |
| Cytokine | Human VEGF | PeproTech | 100-20 |
| Kit | Click-iT TUNEL Alexa Fluor 647 Imaging Assay | ThermoFisher | C10247 |
| Kit | LDH Cytotoxicity Assay Kit | ThermoFisher | C20301 |
| Kit | LEGENDplex Human HSC Panel | BioLegend | 740611 |
| Material | 96-Well No-Bottom Plates | Greiner Bio-One | 655000-06 |
| Material | 96-Well Plate Lids | Greiner Bio-One | 656170 |
| Material | Silicon Wafer | University Wafer | 452 |
| Material | Smooth-Cast 310 | Smooth On | N/A |
| Material | SU-8 2150 | MicroChem | NC0216470 |
| Material | SU-8 Developer | MicroChem | NC9901158 |
| Material | Sylgard 184 Silicone Elastomer Kit | Dow Corning | 2065622 |
| Media | αMEM | Sigma-Aldrich | M0644 |
| Media | EBM-2MV | Lonza | CC-3202 |
| Media | FBS | HyClone | SH3007103 |
| Media | Penicillin-Streptomycin | HyClone | SV30010 |
| Misc. | Biopsy Punches | Integra Miltex | 33-31AA-P/25,<br>33-31-P/25,<br>33-34-P/25 |
| Misc. | nanoDot | Landauer | 03053-OTO |

**Table S2. Antibodies and dilutions.**

| Name | Supplier | Catalog No. | Dilution (Conc.) |
|---|---|---|---|
| Rabbit Anti-CXCL2 | Abcam | ab9797 | 1:100 (5 µg/mL) |
| Rabbit Anti-Fibronectin | Abcam | Ab2413 | 1:100 (10 µg/mL) |
| Rabbit Anti-JAG1 | Abcam | ab7771 | 1:100 (10 µg/mL) |
| Rabbit Anti-Osteopontin | Abcam | ab8448 | 1:100 (82 µg/mL) |
| Rabbit Anti-SCF | Abcam | ab64677 | 1:100 (10 µg/mL) |
| Rabbit IgG Isotype | Abcam | ab37415 | 1:1000 (5 µg/mL) |
| Anti-CD31 AF647 | BioLegend | 303112 | 1:100 (1 µg/mL) |
| Anti-CD34 BV421 | BioLegend | 343609 | 1:100 (1 µg/mL) |
| Anti-CD45 AF488 | BioLegend | 304019 | 1:100 (2 µg/mL) |
| Mouse IgG Isotype BV421 | BioLegend | 400259 | 1:50 (1 µg/mL) |
| Goat Anti-Rabbit AF488 | Life Technologies | A11008 | 1:200 (10 µg/mL) |

**Table S3. Primary cell sources.**

| Name | Supplier | Catalog Number | Lot(s) |
|---|---|---|---|
| BM CD34+ Cells | Lonza | 2M-101 | 0000690910<br>0000573128 |
| HUVEC | Lonza | C2519A | 0000470896 |
| MSCs | RoosterBio | MSC-001 | 00037 |
